# The principles of peptide selection by the transporter associated with antigen processing

**DOI:** 10.1101/2023.06.16.545308

**Authors:** James Lee, Michael L. Oldham, Victor Manon, Jue Chen

## Abstract

The adaptive immune response eliminates infected and cancer cells through the recognition of antigenic peptides displayed by major compatibility complex class I (MHC-I) molecules^1, 2^. A single transporter, the transporter associated with antigen processing (TAP), supplies nearly the entire peptide repertoire for the many MHC-I alleles^3–5^. A fundamental unresolved question is how TAP transports peptides with vast sequence diversity. Here, using cryo-electron microscopy (cryo-EM), we determined seven structures of human TAP in the presence and absence of peptides with different sequences and lengths. We observe that peptides are suspended in the transmembrane cavity of TAP with the peptide N-and C-termini anchored at two distal binding pockets. The central residues of the peptide are unrestricted, making few contacts with TAP. A minimum of eight residues is required to bridge the two binding pockets, aligning with the lower length limit for MHC-I binding^6, 7^. Mutations in TAP that disrupt hydrogen bonds with the peptide termini nearly abolish MHC-I surface expression, indicating that binding depends on interactions with mainchain atoms at the two termini. By utilizing two spatially separated binding pockets and concentrating interactions at the two ends of the peptide, TAP functions as a molecular caliper, selecting peptides for length while permitting sequence diversity.

## Introduction

Cytotoxic T cells recognize and eliminate diseased cells upon the detection of foreign peptides displayed by cell surface MHC-I molecules. The loading of intracellular peptide antigens onto MHC-I molecules occurs in the endoplasmic reticulum (ER) and involves several key steps^1^. In the cytosol, proteins are degraded into small peptides by the proteasome^8, 9^. A small fraction of these peptides is transported into the ER by the transporter associated with antigen processing (TAP)^10–13^. There, high affinity peptides are selectively loaded onto MHC-I by the peptide loading complex and eventually displayed on the surface of the cell for immunosurveillance^2, 14^. Peptides derived from endogenous proteins are immunologically silent due to self-tolerance. In the context of viral infection or malignant transformation, however, displayed peptides derived from foreign or aberrant proteins trigger an immune response that eradicates the diseased cell.

The potential sequence space of foreign peptides is vast, due to the combined diversity of viral proteomes and products of mutational burden in malignant transformation. Display of these peptides thus depends on antigen presentation machinery flexible enough to accommodate broad sequence diversity. The principles governing MHC-I presentation are well understood^6, 15, 16^. Humans have three classical MHC-I (HLA-A,B,C) genes that are highly polymorphic, resulting in vast genetic diversity within a population. Furthermore, by interacting with the peptide’s backbone and a few anchoring residues, each MHC-I allele can present a unique cohort of peptides. In contrast, our understanding of how TAP accommodates vast sequence diversity remains incomplete. Each of us only possesses one TAP, which exhibits very limited degrees of polymorphism^17–21^. One might expect TAP to be promiscuous as this transporter provides peptides to the many MHC-I alleles. However, numerous studies showed that TAP does exercise a certain degree of substrate preference. TAP most efficiently transports peptides of 8-12 residues in length, a size similar to the peptides MHC-I molecules bind^22, 23^. Although the upper size limit is not clearly established, it is widely accepted that the lower limit is 8 residues^22, 24–26^. Also, like MHC-I, free N-and C-termini of the peptide are essential for TAP binding. Modification at either terminus reduces peptide recognition substantially^27–30^. Finally, the sequence tolerance of TAP seems to be broad. It appears that only the N-terminal three positions and the C-terminus determine binding affinity^26, 29, 31, 32^. Previous studies have shown that the peptide is bound in an extended conformation^33^ and that specific residues on TAP influence substrate binding^34–39^. However, molecular details of the TAP-peptide interaction are necessary to understand the functional properties of TAP.

In this study, we used cryo-EM to determine a series of structures of human TAP, both in the absence of peptide and in the presence of peptides with various sequences and lengths. These findings define the molecular basis of how TAP selects peptides of the appropriate length with vast sequence diversity for MHC-I presentation.

## Results

### The structures of TAP in the presence and absence of peptide substrates

Wild-type (WT) human TAP was overexpressed and purified from human embryonic kidney (HEK) cells in the detergent glycol-diosgenin (GDN) (Extended Fig. 1a) for biochemical and structural studies. TAP is a heterodimeric, ATP binding cassette (ABC) transporter consisting of two polypeptide chains, TAP1 and TAP2 (Fig. 1a). Each subunit has three structural domains: an N-terminal transmembrane domain (TMD0) that interacts with other components of the larger peptide loading complex, a six transmembrane helical domain (TMD) that forms the translocation pathway, and a nucleotide binding domain (NBD) that hydrolyzes ATP (Fig. 1a). The TMD0 domains facilitate MHC-I loading, but are not essential for TAP peptide transport activity^40, 41^. The biochemical properties of the recombinant protein were assayed in the presence or absence of a 9-mer peptide containing residues RRYQKSTEL. This peptide has been well-characterized as a substrate transported via TAP^29^ and presented on the cell surface by HLA-B27^42^. Thus, it is designated as the b27 peptide in this work. In the absence of peptide, TAP hydrolyzes ATP at a specific turnover rate of 2 per min (Fig. 1b,c). The b27 peptide stimulates ATP hydrolysis in a dose-dependent manner (Fig. 1b) and, consistent with previous studies^27–30^, stimulation is abolished by blocking either the free amino-or the carboxyl-termini (Fig. 1c).

**Figure 1:**
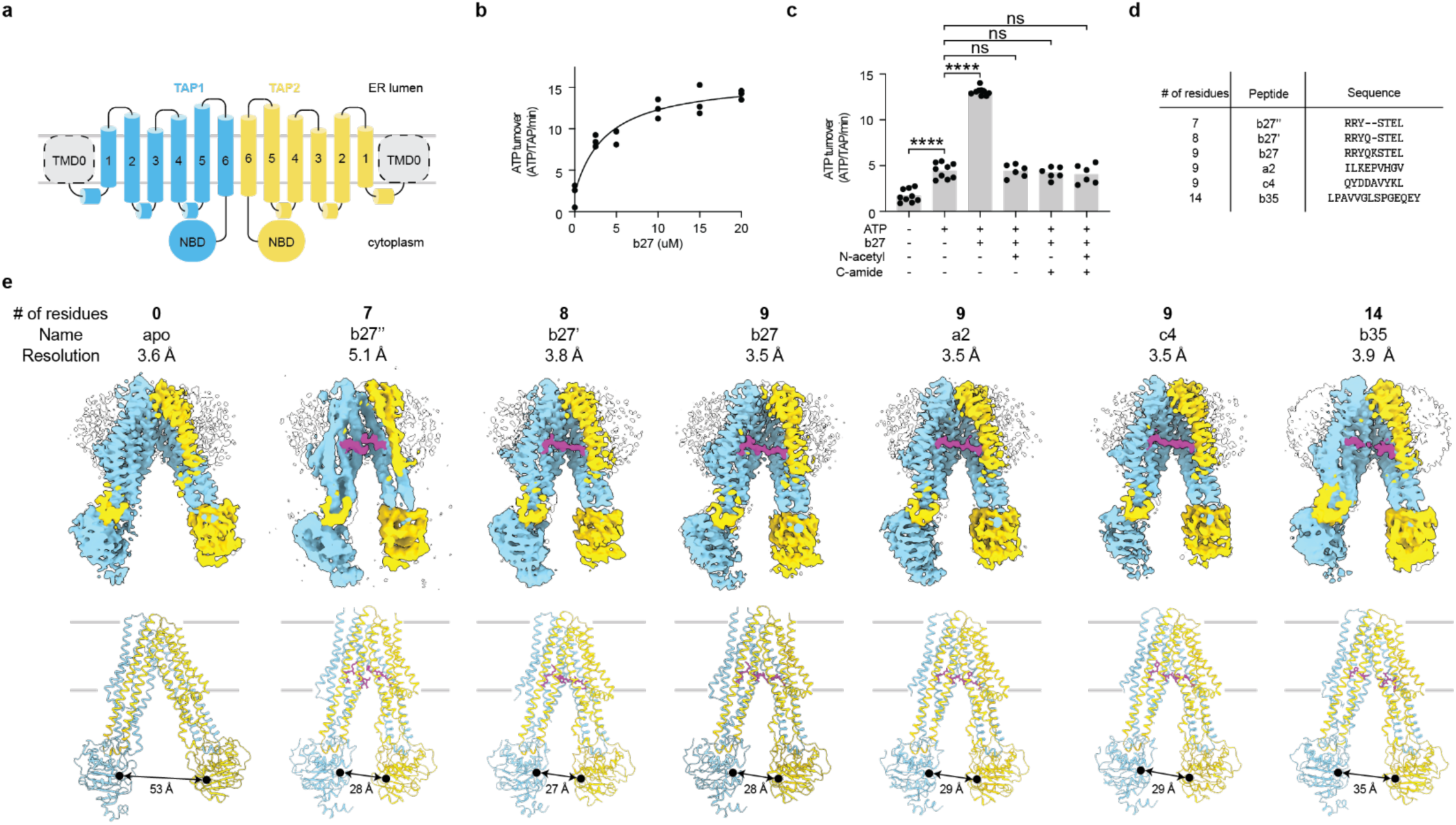
Structures of TAP in the presence and absence of peptide substrates. (a) Topology diagram of the TAP complex. TAP1 and TAP2 are colored in cyan and yellow, respectively. (b) The ATPase activity as a function of b27 concentration. Data represent measurements from 3 technical replicates (n = 3), collected at 28°C with 0.5 µM TAP and 3 mM ATP. (c) ATPase activity in the presence of b27 variants. Data represent the means and standard errors from measurements of 3 technical replicates of 3 biological replicates for the -ATP, +ATP, and +ATP/b27 conditions (n = 9) and 3 technical replicates of 2 biological replicates for the termini modified peptides (n = 6). Concentrations: TAP, 0.5 µM; peptides, 20 µM; ATP, 3 mM. Statistical significance was tested by one-way analysis of variance. NS, not significant. ****, P<0.0001. (d) Peptides used in this study. (e) cryo-EM densities (upper) and ribbon representations (lower) of TAP in the presence and absence of peptide substrates. The density is sliced to show the translocation pathway. TAP1, TAP2, and peptides are colored in cyan, yellow, and violet, respectively. Unassigned densities are colored white. The boundaries of the membrane are marked by grey lines and the distance between TAP1 D668 and TAP2 E632 are indicated.

Using cryo-EM, we determined seven structures of TAP in the absence and presence of peptides ranging from 7 to 14 residues long (Fig. 1d,e, and Extended Fig. 1-3). All structures were determined in the absence of ATP, as previous work showed that ATP reduces the affinity of peptides for TAP^43^. The highest quality maps were obtained with 9-mer peptides, followed by the apo, 8-mer, and 14-mer bound structures. Although the flexibility of the TAP1 NBD impacted the overall resolution, the densities for the TMDs and the bound peptides are well defined, permitting unambiguous assignment of the molecular interactions between TAP and the corresponding peptide substrate (Fig. 1e, 2a, and Extended Fig 3). In the presence of the 7-mer peptide, which is shorter than the preferred minimum length for TAP, the structure exhibits greater heterogeneity, limiting the resolution to 5.1 Å (Extended Fig 1c). In all reconstructions, amorphous densities corresponding to the TMD0 domains are visible only at a lower contour, indicating that the TMD0 of both TAP subunits are flexible.

**Figure 2.**
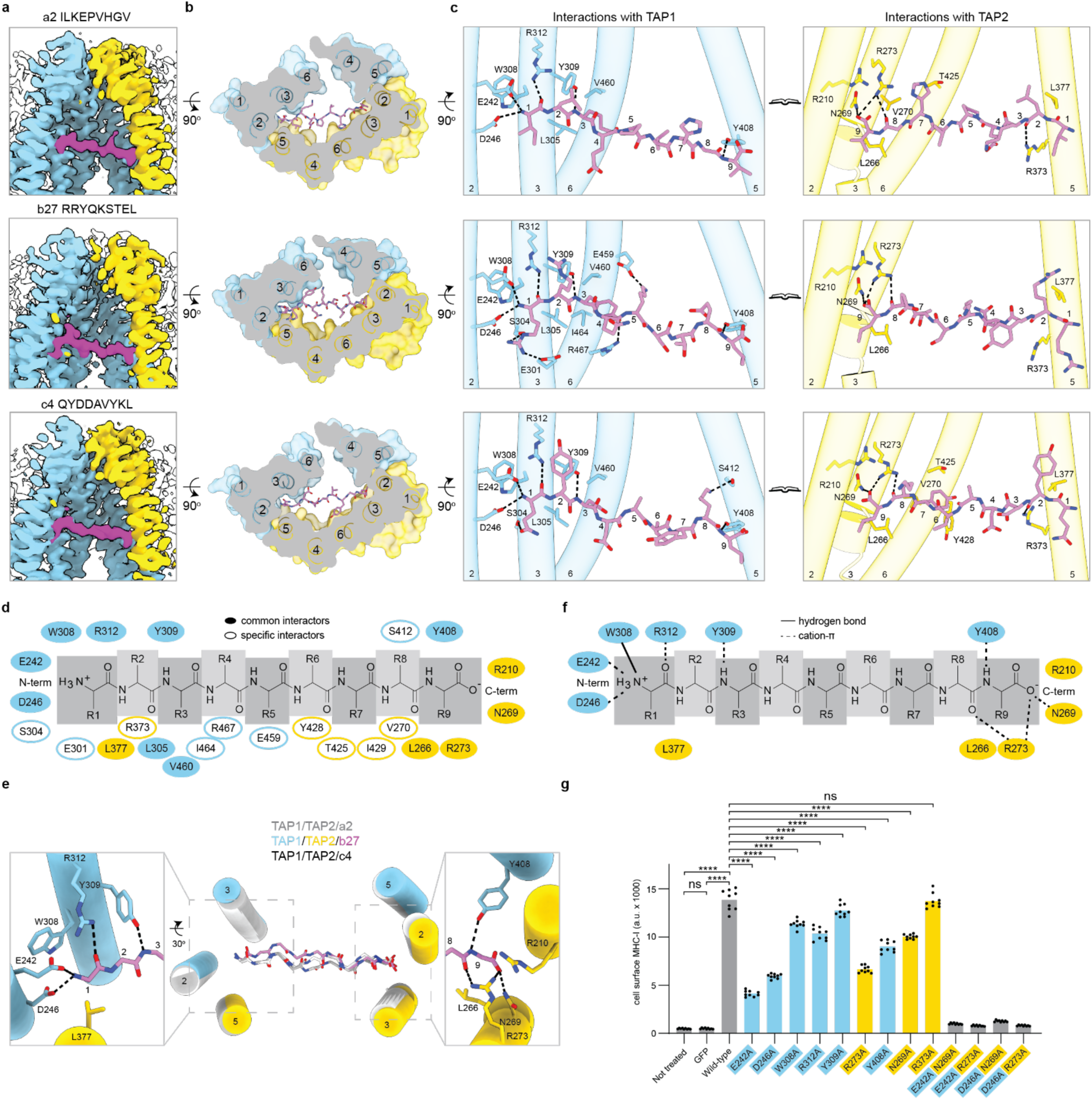
Molecular details of how TAP interacts with 9-mer peptides. (a) cryo-EM density of the TAP translocation pathway. (b) Cross-sectional surface representation of the TAP peptide binding site as viewed from the ER lumen. (c) Peptide-binding residues on TAP1 and TAP2. Only the transmembrane helices that are within 4 Å of the peptide are shown. Hydrogen bonds are marked as black dashed lines. (d) Schematic of the residues in TAP involved in peptide binding. Residues that interact with all three 9-mer peptides are indicated as filled ovals, those that only interact with 1 or 2 peptides are indicated as open ovals. (e) Superposition of the three 9-mer peptides. The side chains are omitted for clarity. Insets display the conserved interactions at the N-and C-termini. (f) Schematic drawing showing the interactions between TAP and the peptide backbone observed for all three 9-mers. (g) Flow cytometry analysis of MHC-I cell surface levels in TAP KO cells expressing GFP-tagged TAP variants. Data represent the median fluorescence intensity of three technical replicates across three biological replicates (n = 9). Statistical significance was tested by one-way analysis of variance. NS, not significant. ****, P<0.0001.

**Figure 3.**
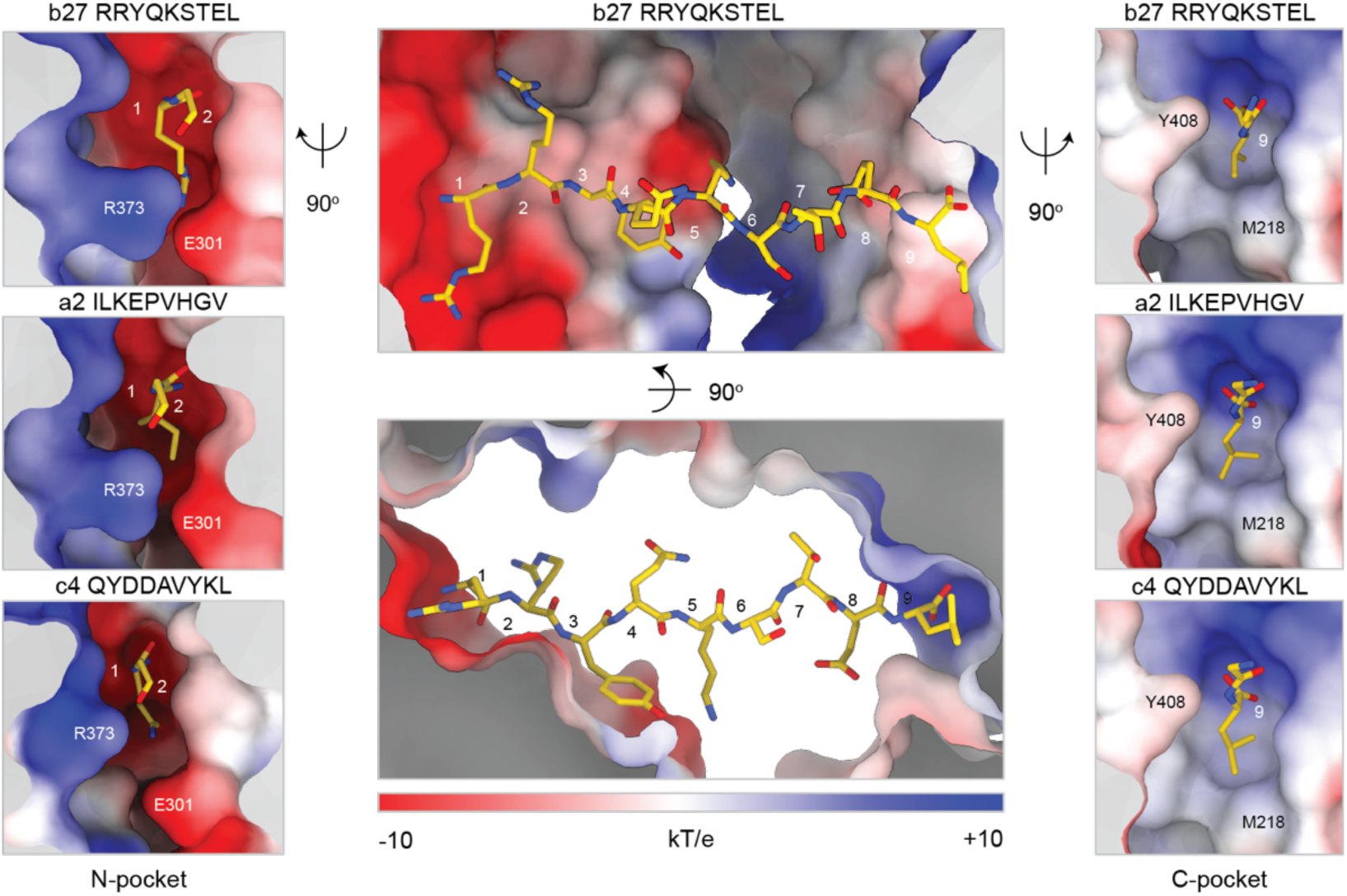
The N-and C-binding pockets. Electrostatic surface representations of the peptide binding site. Center: binding of the entire b27 peptide. Left: insertion of the first residue of the b27 (top), a2 (middle), and c4 (bottom) peptides into the N-pocket. Right: binding of the last residue of each peptide to the C-pocket.

Globally, all the structures exhibit a similar NBD-separated conformation with the peptide-translocation pathway open to the cytoplasm (Fig. 1e). The largest degree of NBD separation is observed in the absence of peptide where the two ATP-binding sites are 53 Å apart. In the presence of peptides, the two halves of TAP move closer towards each other; the most significant change occurs with the 8-mer peptide (26 Å narrower), and the least change is associated with the 14-mer peptide (18 Å narrower).

Two different configurations of the TMDs are observed among the seven structures (Extended Fig. 4). Five structures exhibit a typical inward-facing conformation where the translocation pathway is fully sealed at the ER luminal side (Extended Fig. 4a). However, the structures obtained with the b27 (9-mer) and b27” (7-mer) peptides deviate from this configuration (Extended Fig. 4b-c). Instead, they both display a lateral opening that connects the translocation pathway to the membrane. The gap between the TM helices is filled with lipid molecules, blocking peptide access to the ER lumen (Extended Fig. 4b,d). The occurrence of this lateral opening does not seem to be correlated with the length or sequence of the bound peptide. Furthermore, a similar lateral opening was previously observed in the structure of TAP stabilized by the viral inhibitor ICP47^44, 45^. Whether this opening is an intrinsic step in the peptide transport cycle or merely a reflection of structural flexibility in the absence of other members of the peptide loading complex remains a subject for future investigation.

**Figure 4:**
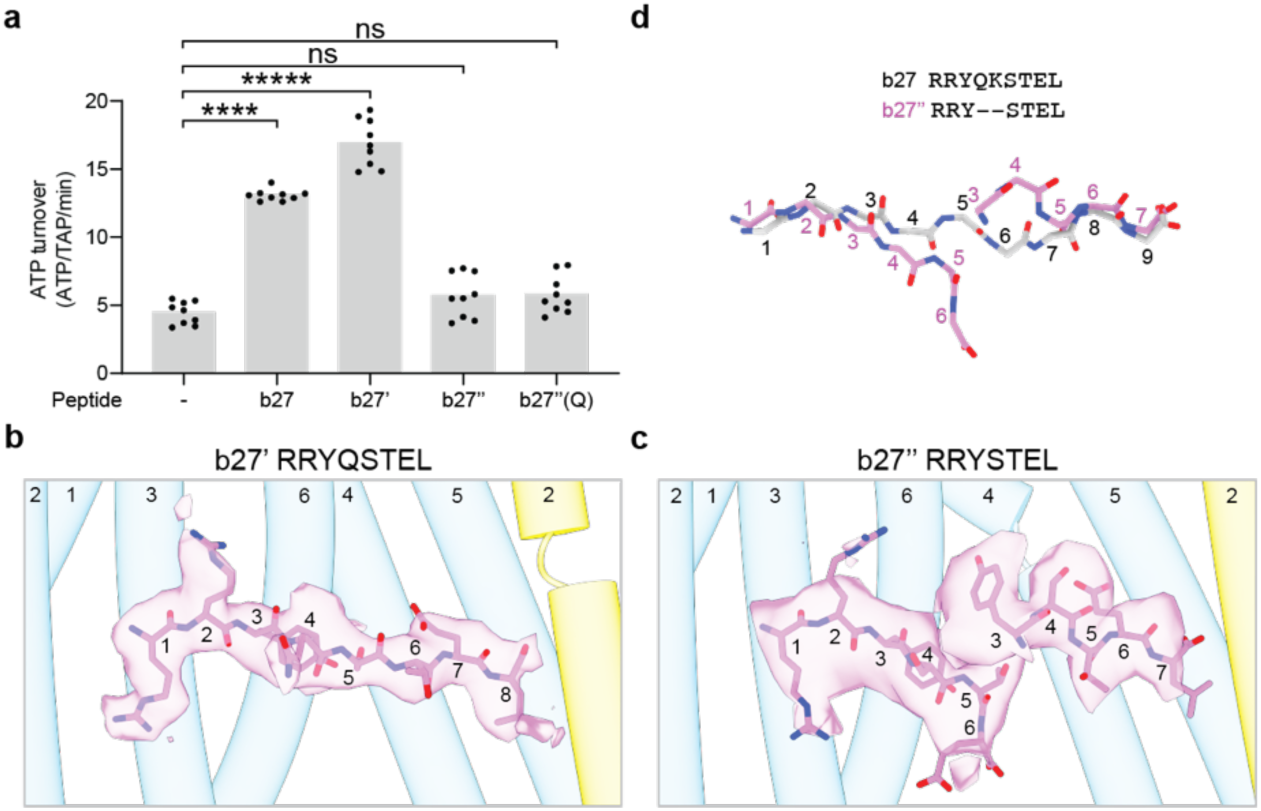
Binding of 8-and 7-mer peptides. (a) The 8-mer (b27’) but not the 7’mer peptides (b27’’ and b27”(Q)) stimulated ATP hydrolysis. Data represent the means and standard errors from measurements of 3 technical replicates of 3 biological replicates (n = 9). Statistical significance was tested by one-way analysis of variance. NS, not significant. ****, P<0.0001. (b-c) EM density for the bound (b) 8-mer b27’ and (c) 7-mer b27’’ peptides, shown as transparent magenta surfaces. (d) Overlay of the peptide backbones of the 9-mer (grey) and the two partially built 7-mer peptides (colored).

### The 9-mer peptides bind in extended conformations parallel to the membrane

To understand how TAP recognizes peptide antigens, we first analyzed the structures of TAP in complex with three 9-mer peptides (Fig. 2), which are optimal in length for binding^22, 23^. These peptides, IL-KEPVHGV (a2), RRYQKSTEL (b27), and QYDDAVYKL(c4), are natural products of the antigen processing pathway being presented by the three representative MHC-I alleles, HLA-A2^46^, HLA-B27^42^, and HLA-C4^47^, respectively.

The structures of the a2-and c4-bound TAP are essentially identical, evident by the overall root mean square deviation (rmsd) of 0.8 Å. In contrast, the b27-bound structure exhibits a different TMDs configuration as described (Extended Fig. 4). In all structures, the peptides are situated in a central cavity encircled by TM 2-6 of both TAP1 and TAP2 (Fig. 2a-b). When viewed from a perspective perpendicular to the membrane plane, each peptide stretches roughly 27 Å across the TM cavity (Fig. 2b). Interactions with TAP are concentrated at the N-and C-terminal ends, primarily involving main chain atoms of the peptide (Fig. 2c-f). Side chains at the first three positions and the last position also variably contact TAP, further enhancing binding in a sequence-dependent manner (Fig 2c-d). The central positions make few contacts, which are exclusively through side chain atoms.

Despite their sequence diversity, the three 9-mer peptides are stabilized in the peptide binding pocket by a common set of interactions involving backbone atoms on separate ends of the peptide (Fig. 2e-f). These interactions, including eight hydrogen bonds, a cation-ρε and two van der Waals’ interactions, enable sequence-independent binding. Specifically, at the N-terminal region, the free amino group of each peptide forms a cation-ρε interaction with W308 and two hydrogen bonds with E242 and D246 in TAP1 (Fig. 2e-f). The carbonyl oxygen and amide at positions 1 and 3 are coordinated through hydrogen bonds by TAP1 R312 and Y309, respectively. Furthermore, TAP2 L377 forms van der Waals’ interactions with the N-terminus of all three peptides and R373 interacts with the main chain atoms in the a2 and c4 peptides but not with the b27 peptide (Fig. 2c-d). In comparison, binding at the C-terminal end of the peptide is dominated by residues from TAP2 (Fig. 2e,f). The terminal carboxyl group of all three peptides are engaged by TAP2 N269 and R273 via hydrogen bonds, as well as by longer-range electrostatic interactions with TAP2 R210. The peptide terminal amide bond is stabilized by van der Waal’s interactions with TAP2 L266 and hydrogen bonds with TAP2 R273 and TAP1 Y408. The latter residue, Y408, was previously identified to interact with the C-terminal end of peptides in mutagenesis studies^18, 48^.

All backbone-binding residues on TAP1 and TAP2 are highly conserved among different species (Extended Fig. 5). To investigate their functional significance in a cellular context, we analyzed the surface expression of MHC-I molecules in cells carrying different TAP variants (Fig. 2g and Extended Fig. 6a-b). In TAP-knockout (KO) cells, MHC-I surface presentation is severely impaired, consistent with the essential role of TAP in providing peptide antigens to complete MHC-I folding. While rescue with wild-type TAP (WT) restores MHC-I surface expression, TAP mutants variably rescue transport depending on the position mutated. Alanine mutations in the N-terminal binding residues, E242, D246, and R273, resulted in the greatest defects in transport with a 50-60% decrease in surface MHC-I level relative to WT. On the other hand, eliminating the hydrogen bond between TAP1 Y309 and the third amide on the peptide resulted in a modest 10% reduction. Mutating R373, which interacts with two out of the three peptides at position 2, had no effect. Most strikingly, combining substitutions at the N-and C-terminal ends reduced MHC-I surface expression to a level similar to that of the TAP KO cells. Loss of function was not due to reduced expression or folding of TAP (Extended Fig. 6c-e). These data indicate that engaging the free amino and carboxyl termini are critical for peptide binding to TAP. This principle holds true in the cellular context, where peptides with variable length and diverse sequences are available for MHC-I presentation.

**Figure 5:**
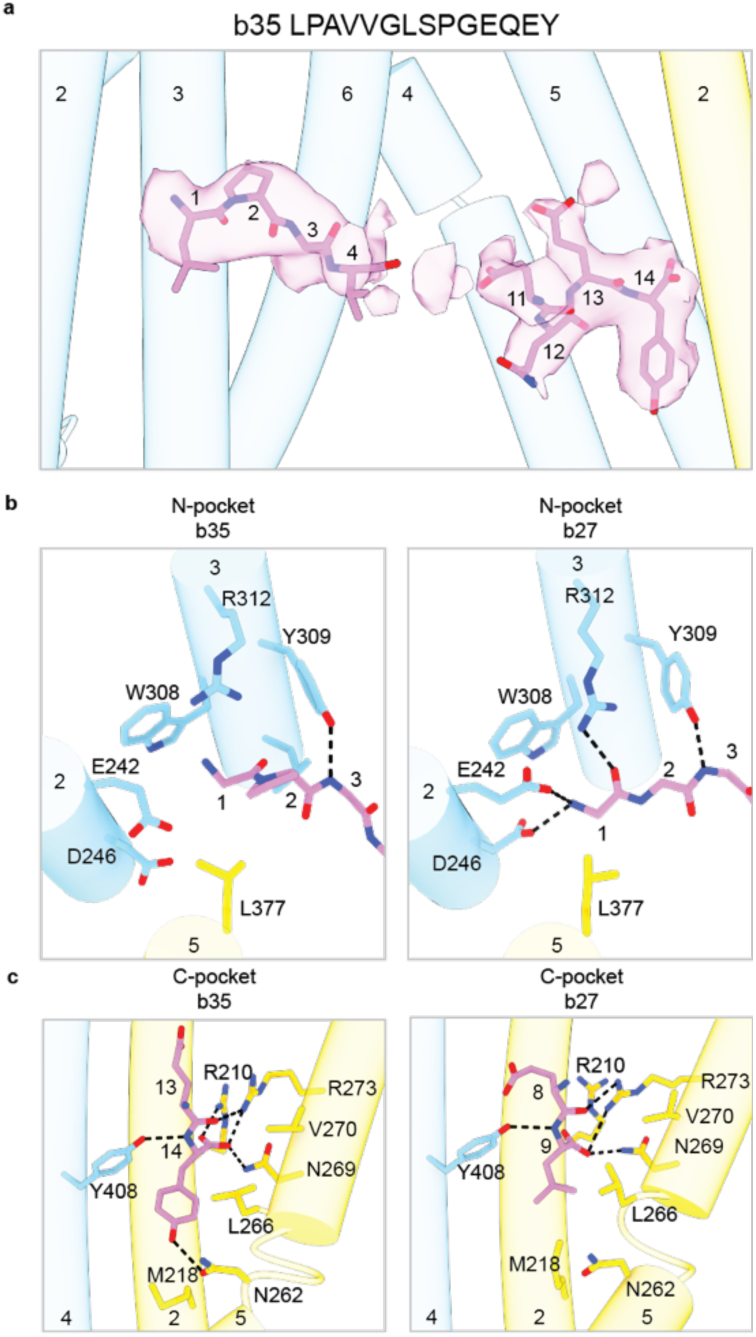
The 14-mer peptide binds through its two termini. (a) Densities corresponding to the 14-mer b35 peptide, shown as transparent magenta surfaces. (b) The b35 peptide makes less interactions with the N-pocket compared to the 9-mer b27. For clarity, only the peptide backbones are shown. (c) At the C-pocket the b35 peptide makes an additional hydrogen bond compared with b27. Only the last two residues of the peptide are shown. The view is in the same orientation as in Fig. 3.

### The N-and C-terminal binding pockets impose sequence preferences

Thus far we have shown that TAP binds peptides primarily through interacting with the peptide backbone, independent of its sequence. Nevertheless, earlier studies demonstrated that TAP does exhibit preferences for specific peptide sequences particularly at the N-and C-termini^29, 31, 32, 49^. Such sequence preferences arise from the structural and electrostatic properties of the peptide-binding site (Fig. 3)

The amino group of each peptide inserts into a deep pocket (the N-pocket) lined with charged and polar residues from TAP1, including E242, D246, W308, and E301 (Fig. 2d and 3). The strong negatively charged surface of the N-pocket and the narrow gate formed by TAP1 E301 and TAP2 R373 explain why the free amino terminus of the peptide is important for high-affinity binding, why a positively charged amino acid at the first position is preferable^29^, and why negatively charged and aromatic residues are disfavored^26, 31^ (Fig. 3).

The C-terminus of the peptide docks into a pocket that differs substantially in size, depth, and electrostatic surface potential from the N-pocket (Fig. 3). The C-pocket can be divided into two regions, a positively charged surface where the terminal carboxyl group binds and a hydrophobic cleft that interacts with the side chain of the C-terminal residue. Previous studies have shown that peptides with a hydrophobic C-terminal residue have a higher affinity for TAP in human, mouse, and the rat cim^u^ alleles^24, 27, 28^. The rat cim^a^ allele is uniquely capable of transporting peptides with a C-terminal basic residue. This functional polymorphism can be attributed to a single amino acid difference in the C-pocket, where all residues are highly conserved except for TAP2 M218 (Extended Fig. 5). In human TAP, M218 lines the hydrophobic region of the C-pocket (Fig. 3, right panels). In the cim^a^ allele, this residue is replaced by a glutamate residue, which is positioned to form ionized hydrogen bond interactions with positively charged side chains at the peptide C-terminus, thus allowing peptides with C-terminal basic residues to be transported.

### A minimum of eight residues are necessary to engage N-and C-pockets concurrently

Many studies have demonstrated that peptides with fewer than eight amino acids are inefficient in competing with model peptides or in being translocated into the ER^24–26^. This length threshold coincides with that of MHC-I molecules, which cannot bind peptides shorter than eight residues^7, 50, 51^. To understand the molecular basis of the minimum length requirement, we modified the b27 peptide to generate shorter peptides and analyzed how they differ from the parent 9-mer (Fig. 4). The 8-mer peptide (b27’) was devoid of the central lysine at position 5 and the two 7-mer peptides each lack an additional residue on either side of the lysine. Compared with b27, the 8-mer peptide increased the ATPase activity of TAP to a slightly greater extent (Fig. 4a). In contrast, neither 7-mer peptides stimulated ATP hydrolysis, suggesting that the compromised function of the 7-mer peptides is likely attributable to their shorter length rather than their sequences.

The structure shows that the 8-mer peptide anchors to TAP at the N-and C-termini through the same interactions as b27 (Fig. 4b and Extended Fig. 7). The overall length of the peptide is shortened by 2 Å, leading to a minor contraction of the translocation pathway (Extended Fig. 7a-b). Intriguingly, the TM helices are arranged similarly to those of the a2-and c4-bound structures (Extended Fig. 7a), rather than that of the b27-bound structure (Extended Fig. 7b). Consequently, the lateral opening seen in the b27 9-mer is no longer present, and the ER luminal gate is entirely closed (Extended Fig. 7d).

In the presence of the 7-mer peptide RRYSTEL (b27”), the conformation of TAP was heterogenous. The final reconstruction, determined at 5 Å resolution, closely resembles that of the b27 9-mer bound conformation (Extended Fig. 1c, 7e), with a clearly visible lateral opening filled by lipid molecules (Extended Fig. 7d). Despite the limited resolution, two distinct densities are apparent within the translocation pathway, each attached to either the N-or C-pockets, respectively (Fig. 4c). The density at the N-pocket is clear enough to model the first six residues of the peptide (Extended Fig. 7f), and the density at the C-pocket can accommodate five residues of the peptide (Extended Fig. 7g). The first three (RRY) and last two residues (EL) of the b27’’ 7-mer peptide within the N-and C-terminal binding sites, respectively, bind in the same way as the b27 peptide (Fig. 4d). The structures of the remainder of the peptide outside of the terminal binding sites diverges. These data suggest that at the concentration of the cryo-EM sample (150 µM), two 7-mer peptides can bind to TAP at the same time. Alternatively, our reconstruction may represent two structures, one with a peptide attached to the N-pocket and the other with a peptide at the C-pocket. In either case, the structural analysis indicates that peptides with less than eight residues cannot span the entire translocation pathway to engage both binding pockets, thus explaining their lower affinity for TAP and inefficiency in competing with longer peptides.

### A longer peptide engages both binding pockets while the central positions are flexible

TAP most efficiently transports peptides 8-12 amino acids in length, but the transport of longer peptides also has been reported^23, 25, 27^. To understand how TAP interacts with longer peptides, we determined the structure of TAP in complex with the 14-mer b35 peptide, LPAVVGLSPGEQEY (Fig. 1e, Fig. 5, Extended Fig. 1e). The b35 peptide contains a proline at position 2, which is supposed to render it a poor substrate for TAP^52, 53^. Nevertheless, this peptide is presented on the cell surface by HLA-B35 in a TAP-dependent manner, suggesting that it is processed through the canonical antigen processing pathway^54^.

In the presence of the 14-mer peptide, the NBDs are separated at a distance intermediate to that of the apo state and 9-mer peptide bound TAP (Fig. 1e). Densities corresponding to the first four and the last four residues of the peptide are clearly observed, but largely absent for the middle six residues (Fig. 5a). Although the b35 peptide engages both binding pockets, important local differences exist compared to how the 9-mer peptides bind TAP (Fig. 5b-c). Specifically, at the N-terminal end, the presence of proline at the second position results in a different backbone conformation that eliminates the hydrogen bonds with D242, E246 and R312 (Fig. 5b). This structural variation is in line with previous findings that prolines are not well-tolerated at position 2^29, 53^. Conversely, the C-terminal end of the b35 peptide makes stronger contacts with TAP compared to the three 9-mer peptides (Fig. 5c). The carboxyl group is situated within 3 Å from R210, forming a salt bridge instead of long-range electrostatic interactions. Additionally, instead of a leucine or valine, the last residue in b35 is a tyrosine, which forms a hydrogen bond with TAP2 N262, a residue previously shown to affect substrate selectivity^38^ (Fig. 5c). This example supports the conclusion from earlier studies that detrimental effects from unfavorable residues can be counterbalanced by favorable residues in other positions^26, 52^.

## Discussion

In this study, we show that TAP holds each peptide at its two ends, leaving the central positions suspended in a large transmembrane cavity. Interactions primarily occur through hydrogen bonds with the backbone atoms of the peptide, although side chains also play a minor role. By concentrating the interactions to the terminal positions and the peptide main-chain atoms, TAP can bind peptides of wide variability in sequence and size. However, a minimum of eight residues in the fully extended conformation is required to bridge the distance between the two binding pockets. Consequently, the lower limit for peptide length is set at eight residues, which aligns with the selection criteria for MHC-I. These principles allow TAP to function as a molecular caliper, selecting peptides of the appropriate length while accommodating vast sequence diversity for MHC-I presentation.

The structural data are consistent with many previous studies. For example, the peptide preferences observed in biochemical studies^26, 29, 31, 32^ can be explained by the configuration of the binding pockets. The distance between the N and C termini of the bound peptide varies in a narrow range between 25-28 Å, comparable to the distances determined by nuclear magnetic resonance and double electron–electron resonance experiments^33, 55^. And finally, many peptide-binding residues, identified through mutagenesis and crosslinking experiments^34–39^, are either in direct contact or near the bound peptide (Extended Fig. 5).

Recently, the structure of a lysosomal peptide transporter TAPL has been determined in multiple conformations^48^. TAPL exhibits approximately 40% sequence identity with TAP, and has broader substrate sequence and length preferences^56, 57^. Although the density of the peptide was inadequate to define molecular interactions, the general position of the bound peptide closely resembles those observed in TAP (Extended Fig 8a-c). However, unlike TAP, TAPL is a homodimer lacking two distinct binding pockets that would enforce a specific N-to C-terminal orientation. Thus, it is possible that the cryo-EM density observed in TAPL reflects a mixture of two orientations of the bound peptide.

Another ABC transporter, the *Thermus thermophilus* multidrug-resistance proteins A and B (TmrAB), has been used in several studies as a model system to understand the molecular mechanism of human TAP^58, 59^. TmrAB and TAP share approximately 30% sequence identity. Cryo-EM analysis of TmrAB revealed density in the transmembrane cavity that was interpreted as the peptide substrate^59^. Structural superposition indicates that this density is proximal to but separate from the peptides bound to TAP (Extended Fig. 8d-f), indicating that TmrAB interacts with its substrate through a mechanism distinct from that of TAP.

Four of the peptides used in this study have also been co-crystallized with MHC-I^42, 46, 47, 54^, providing an opportunity to compare directly their recognition by TAP and MHC-I. Peptides bind to MHC-I in a groove between two α-helices atop a β-sheet. Like TAP, the interactions primarily occur at the two ends of the peptide, involving main chain atoms of the bound peptide. Also similar to TAP, the binding sites in MHC-I for the peptide’s free N-and C-termini are ∼25 Å apart, thereby imposing a minimum size limit of 8 residues for the bound peptide. The major difference from TAP is the involvement of the peptide side chains. MHC-I makes extensive interactions with the side chains of the bound peptide. Certain positions serve as anchors by burying their side chains deep inside binding pockets specific for a particular MHC-I allele. Thus, side chain interactions not only contribute to overall affinity but also dictate allele-specific selectivity in peptide binding^60–63^. In contrast, TAP does not utilize any specific anchoring residues. Favorable side chain interactions can enhance peptide affinity, but unlike MHC-I, they are not essential for binding.

The common and unique structural features of TAP and MHC-I correlate with their functional properties. Given the challenge of achieving both high affinity and promiscuity simultaneously, TAP has evolved to be promiscuous, accommodating greater sequence diversity, while the affinities for peptides can vary over a wide range^26, 29^. In contrast, an individual MHC-I molecule prioritizes affinity to prevent dissociation of the peptide on the cell surface. The diversity of peptide presentation is addressed through genetic polymorphism to ensure protection at the population level. Through this delicate balance, TAP provides a comprehensive peptide reservoir for the numerous MHC-I alleles, while each MHC-I binds a peptide with high affinity to ensure immune fidelity. Such a coordinated mechanism enables us to anticipate peptide antigens from an unpredictable and highly variable pathogenic landscape.

## Acknowledgments

We thank Rui Yan and Zhiheng Yu at the HHMI Janelia Cryo-EM Facility and Mark Ebrahim, Johanna Sotiris, and Honkit Ng at the Evelyn Gruss Lipper Cryo-EM Resource Center at Rockefeller University for assistance in electron microscopy data collection; Svetlana Mazel and Songyan Han at the Flow Cytometry Resource Center at Rockefeller University for assistance in flow cytometry data collection; Kyaw Myo Lwin for the TAP KO cells; and members from the Chen and Mackinnon laboratories for helpful discussions. JL is a Howard Hughes Medical Institute Fellow of the Helen Hay Whitney Foundation and J.C. is an investigator of the Howard Hughes Medical Institute.

## Author Contributions

M.L.O. established the protein purification protocol and collected the first dataset (with the b27 peptide).

J.L. performed all other cryo-EM studies and ATPase assays. J.L. and V.M. carried out the flow cytometry experiments. J.L and J.C. analyzed the data and prepared the manuscript with input from all authors.

## Materials & Correspondence

Cryo-EM maps have been deposited in the Electron Microscopy Data Bank under the accession codes 8T46, 8T4E, 8T4F, 8T4G, 8T4H, 8T4I, and 8T4J. Corresponding atomic models have been deposited in the Protein Data Bank under accession codes EMDB-41021, EMDB-41028, EMDB-41029, EMDB-41030, EMDB-41031, EMDB-41032, and EMDB-41033. All requests for materials should be directed to J.C.

## Methods

### Cell culture

*Spodoptera frugiperda* Sf9 cells (Gibco) were cultured in Sf-900 II SFM medium (Gibco) supplemented with 5% (v/v) heat-inactivated fetal bovine serum (FBS) (Gibco) and 1% (v/v) antibiotic-antimycotic (Gibco) at 27°C. HEK293S GnTI-cells (ATCC) were cultured in Freestyle 293 medium (GIBCO) supplemented with 2% (v/v) FBS at 37°C with 8% CO_2_ and 80% humidity. All cell lines were authenticated by their respective suppliers. Cell lines were tested monthly for mycoplasma contamination by PCR using a Universal Mycoplasma Detection Kit (ATCC) and verified to be negative.

### Protein expression

Human TAP was expressed in HEK293S GnTI-cells based on a previously described protocol^1^. DNA encode for TAP1 with a C-terminal PreScission Protease-cleavable GFP tag and for untagged TAP2 were cloned into separate BacMam vectors to generate pEG TAP1-GFP and pEG TAP2, respectively. To coexpress TAP1 and TAP2, the individual plasmids were combined via SphI and AvrII restriction sites in pEG TAP1-GFP and via SphI and NheI restriction sites in pEG TAP2 to generate pEG TAP1-GFP/TAP2. Bacmid carrying TAP was generated by transforming DH10Bac *E. coli* cells with the pEG TAP1-GFP/TAP2 vector.

Recombinant baculovirus was generated by transfecting Sf9 cells with bacmid using Cellfectin II (ThermoFisher). Baculoviruses were harvested from Sf9 cell media by filtering through a 0.22 µm filter and amplified three times before using for cell transduction. Proteins were expressed in HEK293S GnTI-cells infected with 5% (v/v) of baculovirus at a density of 2.5-3.0 x 10^6^ cells/ml. Cells were induced with 10 mM sodium butyrate 8-12 hours after infection and cultured at 30°C for another 48 hours. Cells were harvested, snap frozen in liquid nitrogen, and stored at −80°C.

### Protein purification

Cells were thawed and resuspended in lysis buffer containing 50 mM HEPES (pH 6.5 with KOH), 400 mM KCl, 2 mM MgCl2, 1mM dithiothreitol (DTT), 20% (v/v) glycerol, 1 μg ml^−1^ pepstatin A, 1 μg ml^−1^ leupeptin, 1 μg ml^−1^ aprotinin, 100 μg ml^−1^ soy trypsin inhibitor, 1 mM benzamidine, 1 mM phenylmethylsulfonyl fluoride (PMSF) and 3 µg ml^−1^ DNase I. Cells were lysed by three passes through a high-pressure homogenizer at 15,000 psi (Emulsiflex-C3; Avestin). Unbroken cells and cell debris were removed by one low speed spin at 4000g for 15 min at 4°C. The supernatant was subjected to a second round of ultracentrifugation at 100,000 x g for 1 hour at 4°C in a Type 45Ti rotor (Beckman) to pellet cell membranes. Membranes were resuspended by manual homogenization in a dounce in lysis buffer supplemented with 1% glycol-diosgenin (GDN) (Anatrace) and incubated for 1 hour at 4°C. The insoluble fraction was removed by centrifugation at 75,000g for 30 min at 4°C and the supernatant was applied to NHS-activated Sepharose 4 Fast Flow resin (GE Healthcare) conjugated with GFP nanobody pre-equilibrated in lysis buffer. After 1 hour, the resin was washed with 10 column volumes of wash buffer containing 50 mM HEPES (pH 6.5 with KOH), 400 mM KCl, 10% glycerol, 1 mM DTT, and 0.01% GDN. To cleave off the GFP tag, PreScission Protease was added to a final concentration of 0.35 mg ml^−1^ and incubated for 12 hours at 4°C. The cleaved protein was eluted with 5 column volumes of wash buffer and collected by passing through a Glutathione Sepharose 4B resin (Cytiva) to remove the PreScission Protease. The eluate was then concentrated using a 15 ml Amicon spin concentrator with a 100-kDa molecular weight cutoff membrane (Millipore) and purified by size exclusion chromatography (SEC) using a Superose 6 Increase 10/300 column (GE Healthcare) pre-equilibrated with SEC buffer containing 50 mM HEPES (pH 6.5 with KOH), 200 mM KCl, 1 mM DTT and 0.004% GDN. Peak fractions were pooled using a 4 ml Amicon spin concentrator with a 100-kDa molecular weight cutoff membrane (Millipore) and used immediately for grid preparation or hydrolysis measurements.

### Cryo-EM grids preparation and data acquisition

TAP purified from gel filtration was concentrated to ∼6 mg ml^−1^ and, where appropriate, incubated with 150 µM of the corresponding peptide on ice for 30 min. Grids were prepared by applying 3.5 µL of this TAP/peptide mixture onto a glow discharged Quantifoil R0.6/1.0 400 mesh holey carbon Au grid with no wait time. The grids were blotted for 3 sec with a blot force of 20 and plunged frozen into liquid ethane using an FEI Mark IV Vitrobot at 6°C and 100% humidity.

All cryo-EM data were collected using a 300 kV Titan Krios transmission electron microscope equipped with a Gatan K3 Summit direct electron detector. All micrographs were collected using SerialEM^2^ in superresolution mode. Data collection parameters for different samples are summarized in Extended Data Table 1.

### Image processing

A similar strategy was employed for cryo-EM data processing for all datasets and a representative flowchart is presented in Extended Fig. 2. Super-resolution image stacks were gain-normalized, binned by 2, and corrected for beam-induced motion using MotionCor2^3^. Contrast transfer functions parameters were estimated using CTFFIND4^4^. Subsequent data processing for all the datasets followed a similar procedure. In general, particles were auto-picked from the motion-corrected micrographs with crYOLO using its general model^5^, extracted in RELION^6^, and imported into cryoSPARC^7^. The picked particles were subjected to multiple rounds of 2D classification, and the resulting particles were subjected to *ab initio* reconstruction with three classes. One class resembled an empty micelle while the other two classes resembled TAP, but with varying continuous density for one of the NBDs. Non-uniform refinement of the best class with the most complete density for the NBDs resulted in a medium-resolution reconstruction of TAP with well-resolved transmembrane helices and NBD2, but with an invisible NBD1. To improve the density of NBD1, all the particles from 2D classification were subjected to iterative rounds of heterogenous refinement using the best reconstruction and a decoy reconstruction with a disordered NBD1 as input models. The resulting particles that gave reconstructions with a complete NBD1 were then subjected to tandem non-uniform refinement followed by local refinement with a protein mask excluding the micelle. These particles were imported into Relion using the csparc2star.py script^8^ and subjected to Bayesian particle polishing^9^ and refined again in cryoSPARC. FSC curves were generated in cryoSPARC, and resolutions were reported based on the 0.143 criterion. Masking and B-factor sharpening were determined automatically in cryoSPARC during refinement.

### Model building and refinement

The sharpened and unsharpened maps from local refinement were used for model building. The initial model was obtained by docking individual domains of the published structure of ICP47-inhibited TAP1/TAP2^10^ into the cryo-EM maps using UCSF ChimeraX^11^. Models were then manually adjusted using Coot where necessary. The models were then iteratively edited and refined in Coot^12^, ISOLDE^13^, and PHENIX^14^. Regions with poor density were removed such as the TMD0 domains or modeled as polyalanine such as the TAP1 NBD. In the case of the TAP2 NBD, a polyalanine homology model based on the crystal structure of mouse TAP1 NBD was first rigid body fit into the density using ChimeraX, manually inspected, and adjusted where necessary. Then, side chains were manually added and refined in Coot. Residues 632-654 and 660-666 in the TAP2 NBDs were modeled as polyalanine. The quality of the final models were evaluated by MolProbity^15^. Refinement statistics are summarized in Extended Data Table 1.

### Analysis of cellular TAP levels

HEK293S GnTI-cells grown in a 6-well plate were infected with 5% P1 baculovirus at 37°C for 36 hours. Cells were harvested by resuspension in 1 ml of buffer containing 50 mM HEPES (pH 8.0 with KOH) and 150 mM KCl and spun down in 1.5 ml tubes for 5 min at 4,000g at 4°C. The cell pellets were resuspended in 1 ml of the same buffer supplemented with 1% GDN and incubated for 60 min at 4°C. Cell lysates were clarified by centrifugation at 20,000g for 2 x 30 min at 4°C. Supernatants were immediately used for analysis by fluorescent size exclusion chromatography (FSEC) or SDS-PAGE. FSEC analyses were performed using a Superose 6 10/300 column (GE Healthcare) pre-equilibrated with SEC buffer. SDS-PAGE analyses were performed at 180V for 75 min using precast 4-20% Tris-HCl polyacrylamide gradient gels (ThermoFisher). In-gel fluorescence was detected using a ChemiDoc MP imaging system (Bio-Rad).

### ATP hydrolysis measurements

Steady-state ATP hydrolysis activity was measured using a nicotinamide adenine dinucleotide (NADH)-coupled assay. Purified TAP was diluted to a final concentration of 0.5 µM in freshly prepared reaction buffer containing 50 mM HEPES (pH 8.0 with KOH), 150 mM KCl, 2 mM MgCl2, 2 mM DTT, 0.004% (w/v) digitonin, 60 µg ml^−1^ pyruvate kinase (Roche), 32 µg ml^−1^ lactate dehydrogenase (Roche), 9 mM phosphoenolpyruvate and 150 µM NADH. Peptide substrates were added to a final concentration of 20 µM unless specified otherwise. 30µl aliquots were dispensed into Corning 384-well Black/Clear Flat Bottom Polystyrene NBS Microplates, and reactions were initiated by the addition of 3 mM ATP. NADH consumption was measured by monitoring the rate of NADH fluorescence depletion at ʎex = 340 nm and ʎem = 445 nm at 28°C using an Infinite M1000 microplate reader (Tecan). Data were converted to ATP turnover with an NADH standard curve. Technical replicates were measured in parallel from the same protein preparation on the same day and biological replicates were measured from different protein preparations from different cells on different days.

### Generation of stable TAP knockout cells

TAP-knockout clonal cells were generated by CRISPR/Cas9-mediated gene editing^16^. gRNAs targeting exon1 of human TAP1 and TAP2 were cloned into the lentiCRISPRv2 vector. The resulting plasmids were transfected into HEK293S GnTI-cells and selected using 1 µg ml^−1^puromycin for 3 days. Puromycin-resistant cells were screened for loss of cell surface MHC-I by flow cytometry and sorted into single cells. After expansion, clones were genotyped by Sanger sequencing.

### Flow cytometry analysis of MHC-I cell surface expression

MHC-I surface expression was analyzed using an allophycocyanin (APC)-coupled antibody W6/32 (eBioscience), which recognizes an epitope shared among all HLA-A,B,C alleles. HEK293S GnTI^-^ cells grown in a 96-well plate were infected with 5% P1 baculovirus encoding wild-type TAP1-GFP/TAP2, a variant, or GFP alone at 37°C for 36 hours. Cells were resuspended in 100 µl FACS blocking buffer (phosphate buffered saline (PBS) supplemented with 5% (w/v) bovine serum albumin (BSA) (Sigma)) and centrifuged at 400g for 5 min at 25°C. Cells were resuspended in FACS blocking buffer supplemented with antibody added at 5 µg ml^−1^ and incubated for 30 min at 4°C in the dark. Subsequently, the cells were washed two times with FACS blocking buffer. The resulting cell pellets were resuspended in FACS buffer (PBS supplemented with 0.5% (w/v) BSA and 0.1% (w/v) sodium azide and counted using an Attune NxT Flow Cytometer (ThermoFisher). Gating for live cells with moderate levels of the GFP were used to compare MHC-I expression. Data were analyzed using FCS Express (De Novo Software). Technical replicates were measured in parallel from different cells on the same day and biological replicates were measured from different cells on different days.

### Data quantification and statistical analysis

All the details of the data and statistical analysis can be found in the figure legends and methods. All data values are presented as mean and standard errors. Statistical significance was assessed by a one-way analysis of variance relative to the wild-type using Graphpad Prism 9.

### Figure preparation

Cryo-EM maps and atomic models depictions were generated using UCSF ChimeraX^11^. Maps colored by local resolution were generated using cryoSPARC. Multiple sequence alignments were generated using Clustal Omega^17^. Surface interface analysis was performed using PDBePISA^18^. Graphs and associated statistics were prepared using GraphPad Prism 9. Software used in this project was managed by SBGrid^19^. All figures were prepared using Adobe Illustrator.

---

**Extended Data Table 1:** Cryo-EM data collection, refinement, and validation statistics

|  | Apo TAP | TAP<br>a2 peptide | TAP<br>b27 peptide | TAP<br>c4 peptide |
| --- | --- | --- | --- | --- |
|  | EMDB-41021<br>PDB 8T46 | EMDB-41028<br>PDB 8T4E | EMDB-41029<br>PDB 8T4F | EMDB-41030<br>PDB 8T4G |
| **Data collection and processing** |  |  |  |  |
| Magnification | 105,000 | 105,000 | 105,000 | 105,000 |
| Voltage (kV) | 300 | 300 | 300 | 300 |
| Electron exposure (e<sup>-</sup>/Å<sup>2</sup>) | 50 | 50 | 66 | 50 |
| Defocus range (μm) | 0.8 to 2.5 | 0.8 to 2.5 | 0.8 to 2.5 | 0.8 to 2.5 |
| Pixel size (Å) | 0.839 | 0.839 | 0.676 | 0.676 |
| Symmetry imposed | C1 | C1 | C1 | C1 |
| Initial particle images (no.) | 3,395,985 | 2,165,210 | 1,618,719 | 2,214,564 |
| Final particle images (no.) | 79,724 | 95,683 | 73,744 | 48,686 |
| Map resolution (Å) | 3.6 | 3.5 | 3.5 | 3.5 |
| FSC threshold | 0.143 | 0.143 | 0.143 | 0.143 |
| **Refinement** |  |  |  |  |
| Map sharpening *B* factor (Å<sup>2</sup>) | 111.6 | 82.5 | 83.6 | 70.6 |
| Model composition |  |  |  |  |
| Non-hydrogen atoms | 7881 | 7714 | 7794 | 7804 |
| Protein residues | 1113 | 1089 | 1079 | 1108 |
| *B* factors (Å<sup>2</sup>) |  |  |  |  |
| Protein | 68.82 | 47.26 | 47.11 | 69.83 |
| R.m.s. deviations |  |  |  |  |
| Bond lengths (Å) | 0.003 | 0.003 | 0.003 | 0.003 |
| Bond angles (°) | 0.550 | 0.553 | 0.571 | 0.534 |
| Validation |  |  |  |  |
| MolProbity score | 1.31 | 1.42 | 1.12 | 1.34 |
| Clashscore | 5.48 | 7.72 | 3.32 | 5.37 |
| Poor rotamers (%) | 0.84 | 0.28 | 0.56 | 0.14 |
| Ramachandran plot |  |  |  |  |
| Favored (%) | 96.92 | 98.05 | 98.34 | 97.78 |
| Allowed (%) | 2.08 | 1.85 | 1.66 | 2.22 |
| Disallowed (%) | 0.00 | 0.09 | 0.00 | 0.00 |

|  | TAP<br>b27'' peptide<br>EMDB-41032<br>PDB 8T4I | TAP<br>b27' peptide<br>EMDB-41031<br>PDB 8T4H | TAP<br>B35 peptide<br>EMDB-41033<br>PDB 8T4J |
| --- | --- | --- | --- |
| **Data collection and processing** |  |  |  |
| Magnification | 105,000 | 105,000 | 105,000 |
| Voltage (kV) | 300 | 300 | 300 |
| Electron exposure (e<sup>-</sup>/Å<sup>2</sup>) | 50 | 50 | 66 |
| Defocus range (μm) | 0.8 to 2.5 | 0.8 to 2.5 | 0.8 to 2.5 |
| Pixel size (Å) | 0.839 | 0.839 | 0.676 |
| Symmetry imposed | C1 | C1 | C1 |
| Initial particle images (no.) | 1,849,805 | 2,301,048 | 1,317,654 |
| Final particle images (no.) | 24,447 | 35,603 | 69,408 |
| Map resolution (Å) | 5.1 | 3.8 | 3.9 |
| FSC threshold | 0.143 | 0.143 | 0.143 |
| **Refinement** |  |  |  |
| Map sharpening *B* factor (Å<sup>2</sup>) | 418.2 | 73.0 | 59.6 |
| Model composition |  |  |  |
| Non-hydrogen atoms | 7811 | 7804 | 7804 |
| Protein residues | 1099 | 1108 | 1108 |
| *B* factors (Å<sup>2</sup>) |  |  |  |
| Protein | 47.72 | 79.3 | 99.2 |
| R.m.s. deviations |  |  |  |
| Bond lengths (Å) | 0.004 | 0.004 | 0.003 |
| Bond angles (°) | 0.610 | 0.659 | 0.585 |
| Validation |  |  |  |
| MolProbity score | 1.13 | 1.49 | 1.49 |
| Clashscore | 3.38 | 6.46 | 8.69 |
| Poor rotamers (%) | 0.84 | 0.43 | 0.14 |
| Ramachandran plot |  |  |  |
| Favored (%) | 98.34 | 97.30 | 97.9 |
| Allowed (%) | 1.66 | 2.60 | 2.10 |
| Disallowed (%) | 0.00 | 0.09 | 0.00 |

**Extended Figure 1:**
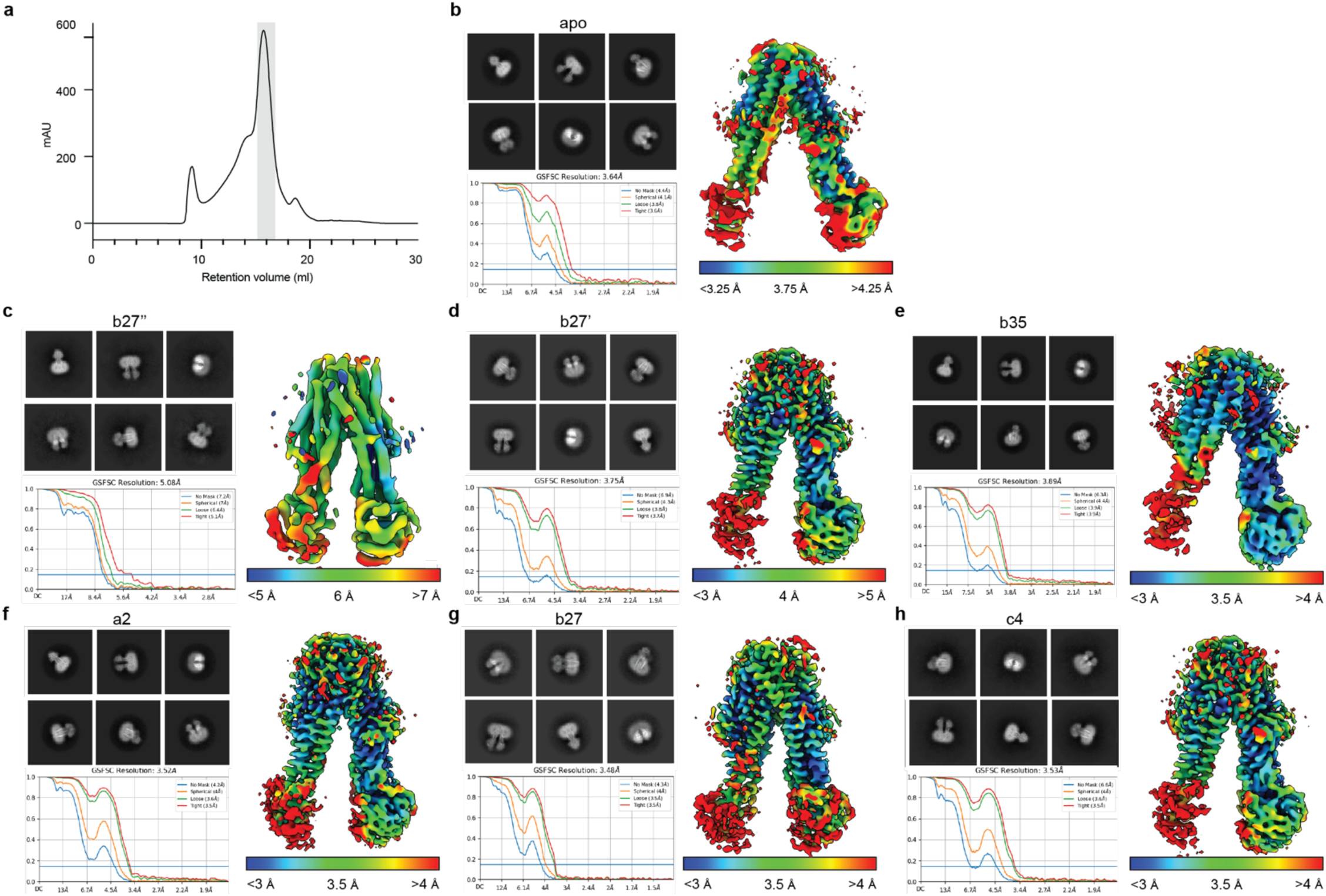
Summary of structural determination. (a) Representative size exclusion profile of purified TAP. Fractions collected for cryo-EM and ATPase assays are marked in grey. (b-h) Representative 2D classes, the Fourier shell correlation (FSC) curve, and colored local resolution maps of the seven structures determined in this study.

**Extended Figure 2:**
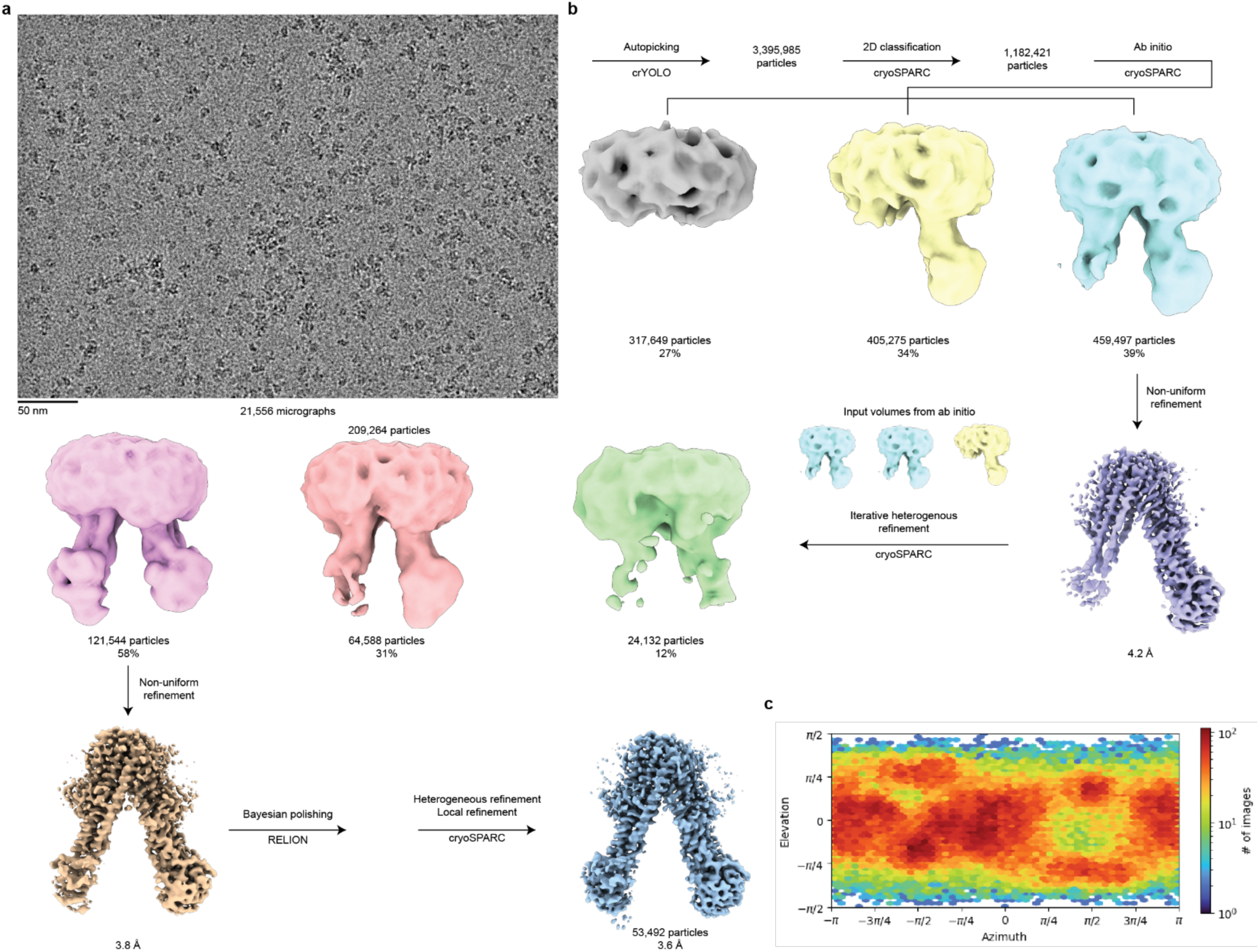
A representative flow chart for cryo-EM data processing of apo TAP. (a) Representative micrograph. (b) Summary of data processing steps. (c) Angular distribution plot for the final map from cryoSPARC.

**Extended Figure 3:**
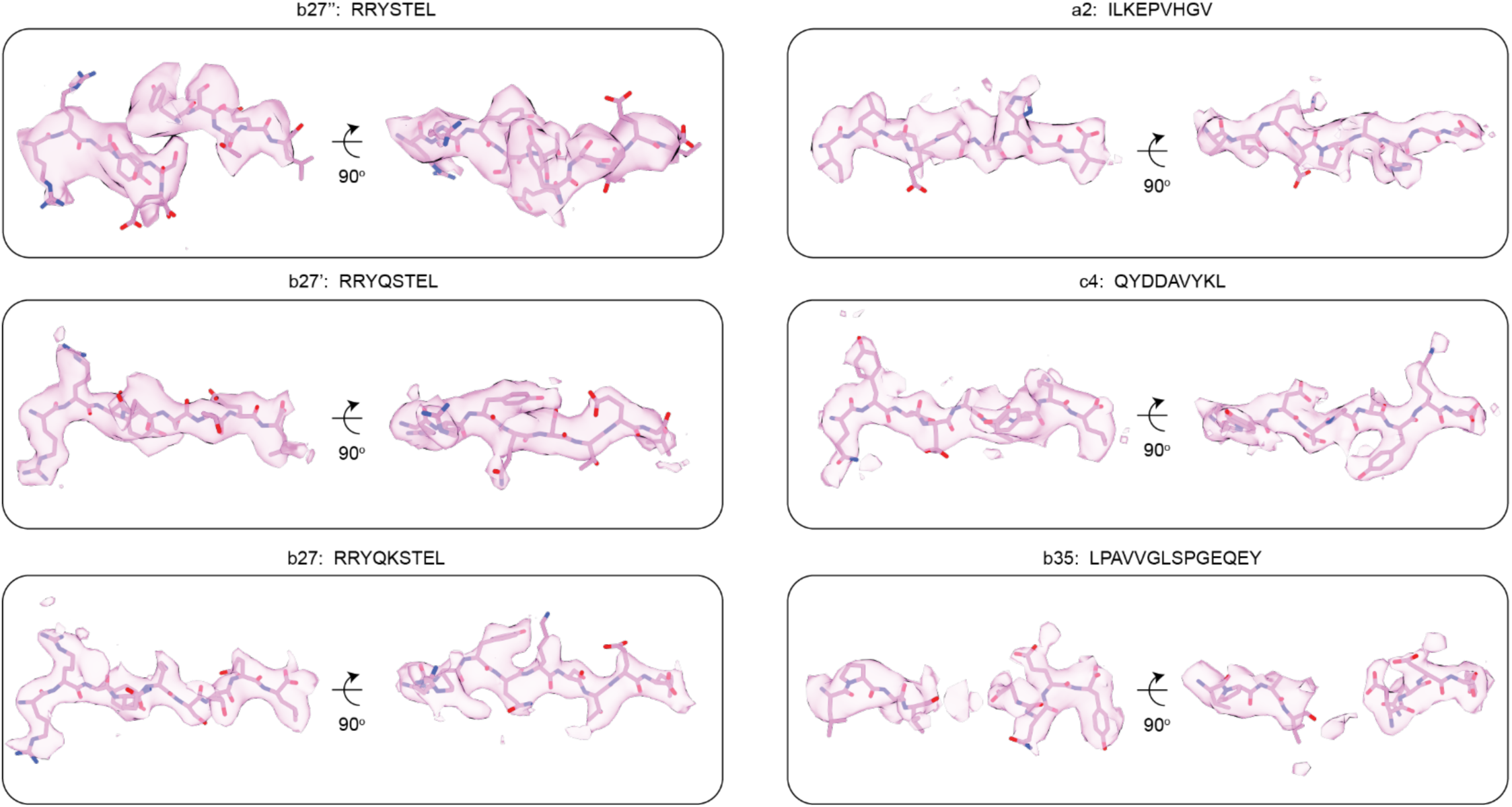
Two views of the cryo-EM density corresponding to each peptide substrate.

**Extended Figure 4:**
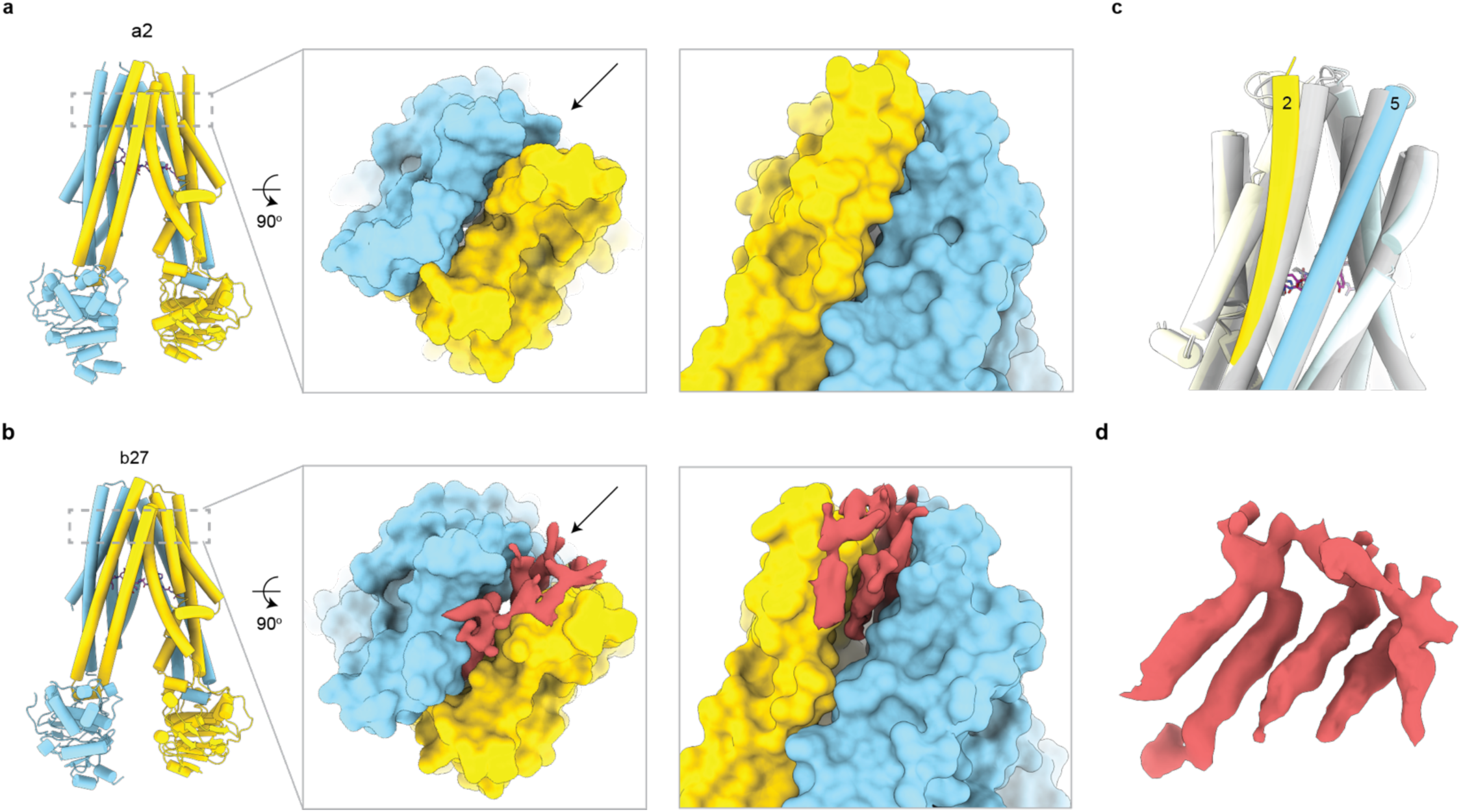
Two different TMD configurations. (a) The structure of a2-bound TAP exhibits a typical inward-facing conformation where the luminal gate (marked by an arrow) is closed. (b) The b27-bound TAP opens to the ER lumen through a lateral gate. (c) Superposition of the structures of a2 (silver)-and b27 (colored)-bound TAP. (d) Densities corresponding to lipids (red) fill the luminal gate in the b27-bound structure.

**Extended Figure 5:**
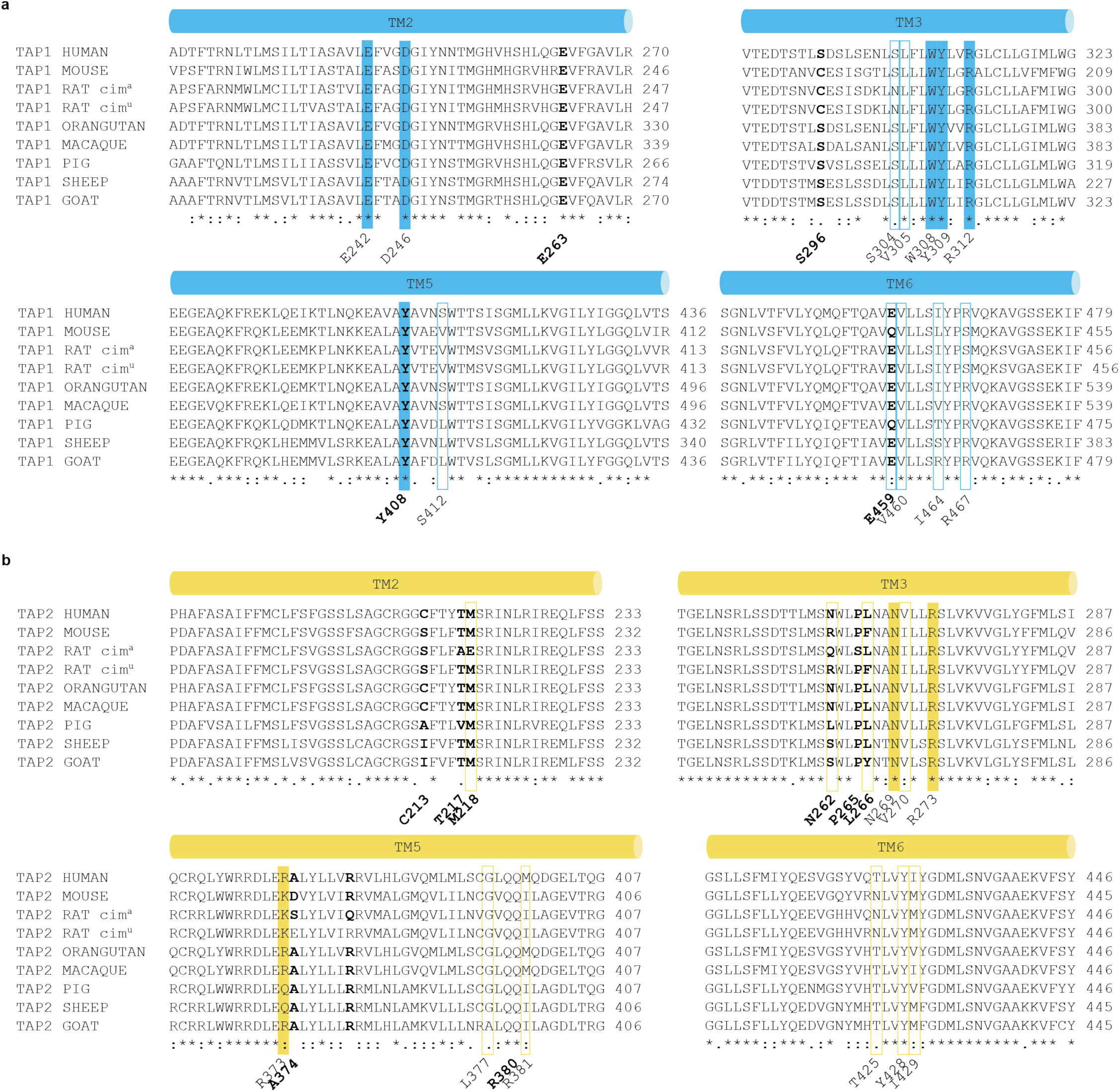
Sequence alignment of. (a) TAP1 and (b) TAP2. Residues identified in this study implicated in peptide binding with either the peptide backbone or peptide side chains are in solid color or outlined, respectively. Residues previously implicated in peptide binding are shown in bold.

**Extended Figure 6:**
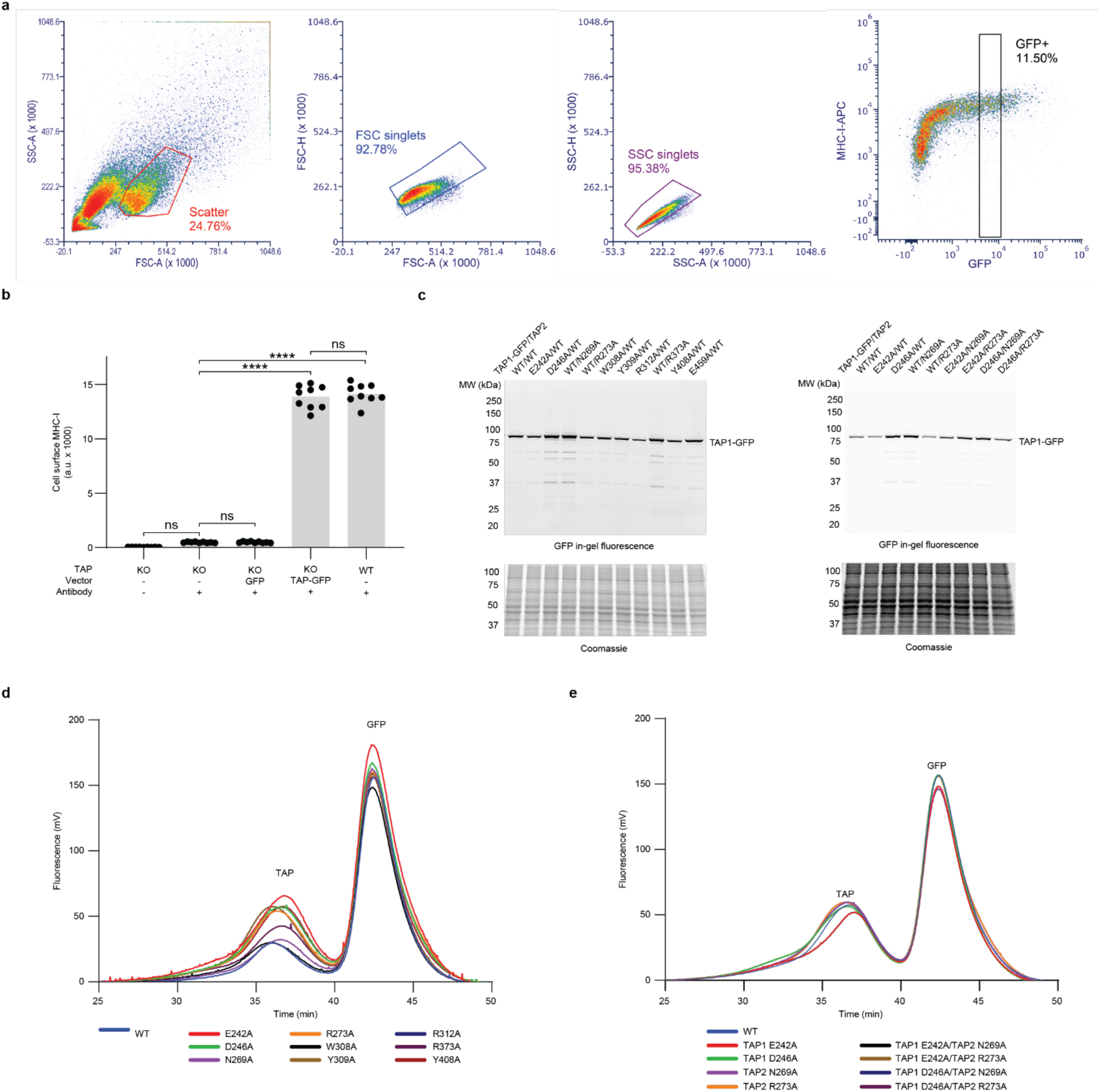
Characterization of the TAP binding site variants. (a) Representative gating strategy for flow cytometry analysis. Single cells were gated by forward and side scatter. (b) Flow cytometry analysis of the TAP KO cell lines. Data represents the means and standard errors from measurements of 3 technical replicates of 3 biological replicates (n = 9). KO, KO GFP, and KO TAP-GFP samples are reproduced from Fig. 2g. (c) SDS-PAGE of lysates of cells expressing different GFP-tagged TAP variants visualized by (top) in-gel fluorescence and (bottom) Coomassie stain. (d-e) Fluorescence size exclusion chromatography from lysates of cells expressing different GFP-tagged variants. WT, TAP1 E242A, TAP1 D246A, TAP2 N269A, TAP2 R273A samples in (e) are reproduced from (d).

**Extended Figure 7:**
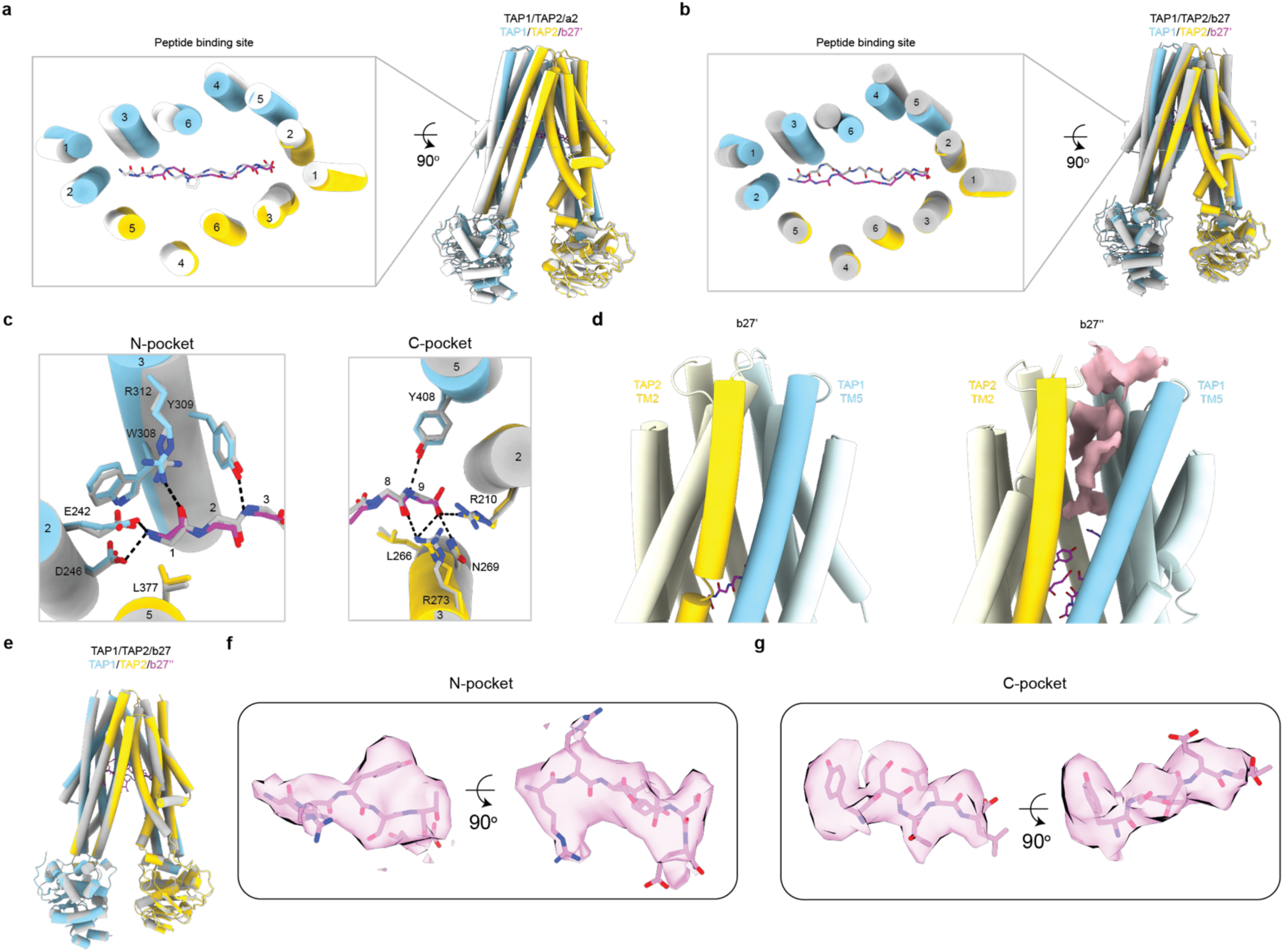
Structural features in the presence of shorter peptides. (a-b) Superposition of the 8-mer b27’-bound structure with the (a) a2-or (b) b27-bound structures. (c) Comparison of local conformation of the b27-and b27’-bound structures. The N-pocket (left) is aligned to TM2 and TM3 of TAP1 and the C-pocket (right) is aligned to TM2 and TM3 of TAP2. (d) The luminal gate is closed in the b27’-bound structure, but open in the b27’’-bound conformation. (e) Superposition of the 9-mer and 7-mer-bound structures. (f-g) Cryo-EM density corresponding to the N-terminally (f) and C-terminally (g) bound 7-mer peptides.

**Extended Figure 8:**
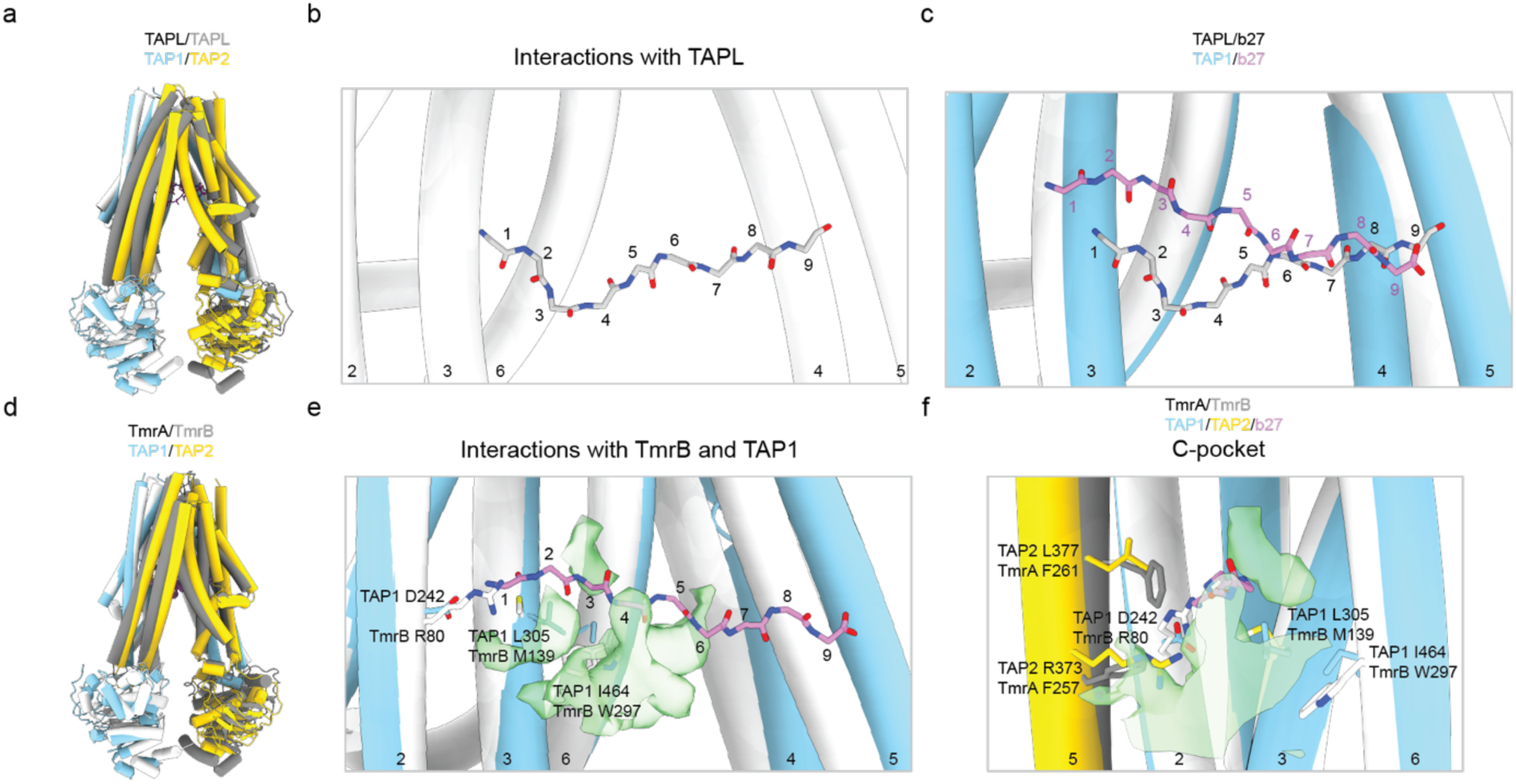
Structural comparisons with TAPL and TmrAB. (a) The overall superposition of b27-bound TAP with b27-bound TAPL (PDB 7VFI) (b) The TAPL peptide-binding site. (c) Comparison of the TAP and TAPL peptide binding sites. Only the mainchain atoms of the b27 peptide are shown. (d) The overall superposition of b27-bound TAP with the inward-facing wide conformation of TmrAB (PDB 6RAN) (e-f) Superposition of the peptide-binding sites of TAP (blue/yellow) and TmrAB (grey). The TAP-bound b27 peptide is shown as magenta sticks and the peptide density in TmrAB is shown in green (EMD-4781). Residues near the density are shown as sticks.

